# Natural constraints explain working memory capacity limitations in sensory-cognitive models

**DOI:** 10.1101/2023.03.30.534982

**Authors:** Yudi Xie, Yu Duan, Aohua Cheng, Pengcen Jiang, Christopher J. Cueva, Guangyu Robert Yang

## Abstract

The limited capacity of the brain to retain information in working memory has been well-known and studied for decades, yet the root of this limitation remains unclear. Here we built sensory-cognitive neural network models of working memory that perform tasks using raw visual stimuli. Contrary to intuitions that working memory capacity limitation stems from memory or cognitive constraints, we found that pre-training the sensory region of our models with natural images imposes sufficient constraints on models to exhibit a wide range of human-like behaviors in visual working memory tasks designed to probe capacity. Examining the neural mechanisms in our model reveals that capacity limitation mainly arises in a bottom-up manner. Our models offer a principled and functionally grounded explanation for the working memory capacity limitation without parameter fitting to behavioral data or much hyperparameter tuning. This work highlights the importance of developing models with realistic sensory processing even when investigating memory and other high-level cognitive phenomena.

## 1 Introduction

Working memory allows us to temporarily retain information and make complex decisions beyond reflexive responses to stimuli [1]. One enduring puzzle in cognitive neuroscience is the seemingly limited capacity of working memory [2, 3]. For instance, in a classical visual working memory paradigm [4], a subject is shown a set of colored squares, and after a brief delay, presented with another set that is either identical or has one of the squares with changed color. Humans’ or other animals’ performance in detecting the color change declines markedly as the number of colored squares (set size) increases – a phenomenon known as the set size effect. It remains perplexing that our ability to perform this straightforward task is so restricted, despite the potential involvement of billions of neurons in the brain.

Numerous studies over the past decades have led to phenomenological models that accurately fit and predict human behavior in several dominant (visual) working memory paradigms [4–10]. Yet, the fundamental question about the origin of the working memory capacity limitation remains a puzzle. We propose that a comprehensive theory of working memory capacity limitation should satisfy the following three desiderata. It should (1) account for the wide range of behavioral patterns of humans and animals in working memory tasks used to study capacity, (2) explain the neural mechanisms underlying these limitation, and (3) offer a principled or functionally grounded explanation for the observed limitation. In this study, we aim to develop models that fulfill all three desiderata, a feat that existing models generally do not satisfy.

Two main classes of phenomenological models, discrete slot [4, 5, 7, 11] and continuous resource [6, 8, 12] models, have been proposed to quantitatively explain human behavior in working memory tasks used to probe capacity, particularly in the visual domain. Slot models posit a limited number of discrete memory slots to hold stimuli. “items” that are allocated slots will be remembered with high fidelity, while information about stimuli without a slot is lost [7]. In contrast, resource models posit that individuals possess continuous resources that can be flexibly divided among items [8]. The total amount of resources is limited and can be allocated to encode multiple stimuli, resulting in varying degrees of memory fidelity. In addition, signal detection models explain the set size effect by the decreasing signal-to-noise ratio when more stimuli are in memory [9, 13]. Although accounting for detailed behavioral measurements, these models are usually descriptive and do not give a principled answer to why we have a particular capacity limitation. The number of slots, amount of resources, or noise are fit to the behavioral data instead of being derived from a principled account. Furthermore, it is hard to map the abstract psychological constructs in these models to neural mechanisms. More recent models try to give a more normative account for the limitation in terms of the trade-off between encoding precision and neural spiking cost [10, 14, 15]. However, assumptions in these models, such as neural cost, are hard to measure and validate in experiments.

Several neural mechanistic theories have been developed to explain the capacity limitation of working memory [16–19]. For example, ring attractor models propose that storing multiple items leads to interference between the attractors representing each item, thereby limiting capacity [16]. Another class of theories posits that multiple objects are encoded at different oscillation phases, and only a limited number of items are encoded at each cycle [18, 20]. However, the extent to which these neural mechanistic models can explain a wide range of behavioral data in capacity-probing tasks remains to be determined.

Functional accounts for the working memory capacity limitation have also been put forth. These theories generally argue that the capacity limitation is either functionally advantageous [21, 22] or a byproduct of another function, such as flexibility [17, 23] or predicting the future [23]. In addition, research in child development suggests that capacity limitation may be beneficial for language acquisition [24]. However, how these insights could be applied to build models of working memory to explain behavioral and neural data is mostly unexplored.

To bridge the gap between varying perspectives on working memory capacity limitations, it is crucial to examine the fundamental units used for measuring capacity. Working memory capacity is often measured in basic units of “items” [3], and resources or slots are also allocated to “items” [7, 8]. However, the definition of an “item” in real-world scenarios remains unclear. Previous models of working memory capacity often assume that stimuli are already disentangled and encoded into individual features or items, with little explanation of how this process occurs. Understanding working memory limitations necessitates a deeper exploration of realistic sensory processing, which previous models have largely overlooked. Previous models generally fail to account for the potential impact of realistic sensory processing, as they often use simplified sensory representations as inputs to the memory system. Moreover, these oversimplified assumptions restrict the applicability of these models across a broad range of tasks since each new task requires the creation of new encoding assumptions. Furthermore, these assumptions are inapplicable to working memory tasks involving realistic objects [25].

To investigate the long-standing puzzle of working memory capacity limitations, we bypass the assumptions of basic units (“items”) of working memory by constructing models that perform working memory tasks using raw sensory inputs in pixel space. Capitalizing on advancements in modeling the ventral visual stream [26–28] and frontoparietal regions [29, 30], we develop sensory-cognitive neural network models that combine a visual system and a cognitive system. Distinct from previous models of working memory, our approach introduces a new class of models without explicit assumptions of limited slots, resources, or noise. In our models, we found that the combination of two parsimonious design principles - naturalistic sensory constraints and functional training objectives of working memory tasks - produces models that match a broad spectrum of behavioral data related to capacity limitations. Our model presents a normative and functionally grounded explanation that elucidates both the behavioral and neural aspects of visual working memory, fulfilling all three previously mentioned model desiderata.

## 2 Results

### 2.1 Image-computable sensory-cognitive models of visual working memory

We developed neural network models that perform visual working memory tasks closely resembling those commonly used in human and animal experiments, using raw visual stimuli in pixel space (Fig.1a). Each model consists of two primary components: the sensory system and the cognitive system (Fig.1b). The sensory system, which emulates the ventral visual stream, is modeled using a convolutional neural network (CNN) [26–28]. The cognitive system, representing the prefrontal and/or parietal regions, is modeled using a recurrent neural network (RNN). RNNs are widely used to model neural dynamics and geometry in cognitive regions of the brain [29, 30, 32]. The CNN processes visual inputs, while the RNN supports short-term memory throughout the delay period. Additionally, the RNN generates top-down attentional modulation to the CNN. This attentional modulation enhances model performance; however, removing it does not alter the central claims of this paper.

In a single trial, the model sequentially receives a series of images, including blank images for the delay period, and generates a sequence of behavioral outputs. At each time step in a trial, the image is input into the CNN, and its output is subsequently fed into the RNN, along with the previous RNN hidden state. The RNN sends top-down feedback to modulate the CNN processing at the next time step. The behavioral output is derived from the RNN’s hidden units (Fig.1c). Depending on the specific task, the output may be a binary decision or continuous variables indicating saccades or other movements.

Various training schemes for the models can be used. One approach involves training the entire sensory-cognitive model “end-to-end” using supervised learning to perform a specific working memory task. Alternatively, the sensory region could be pre-trained on different tasks using natural images [33, 34], including image classification tasks [35] using labeled images or (contrastive) self-supervised learning tasks [36] without using labels. After this pre-training, the sensory region is frozen, and the rest of the model is trained to perform the cognitive task using supervised learning.

**Fig. 1.**
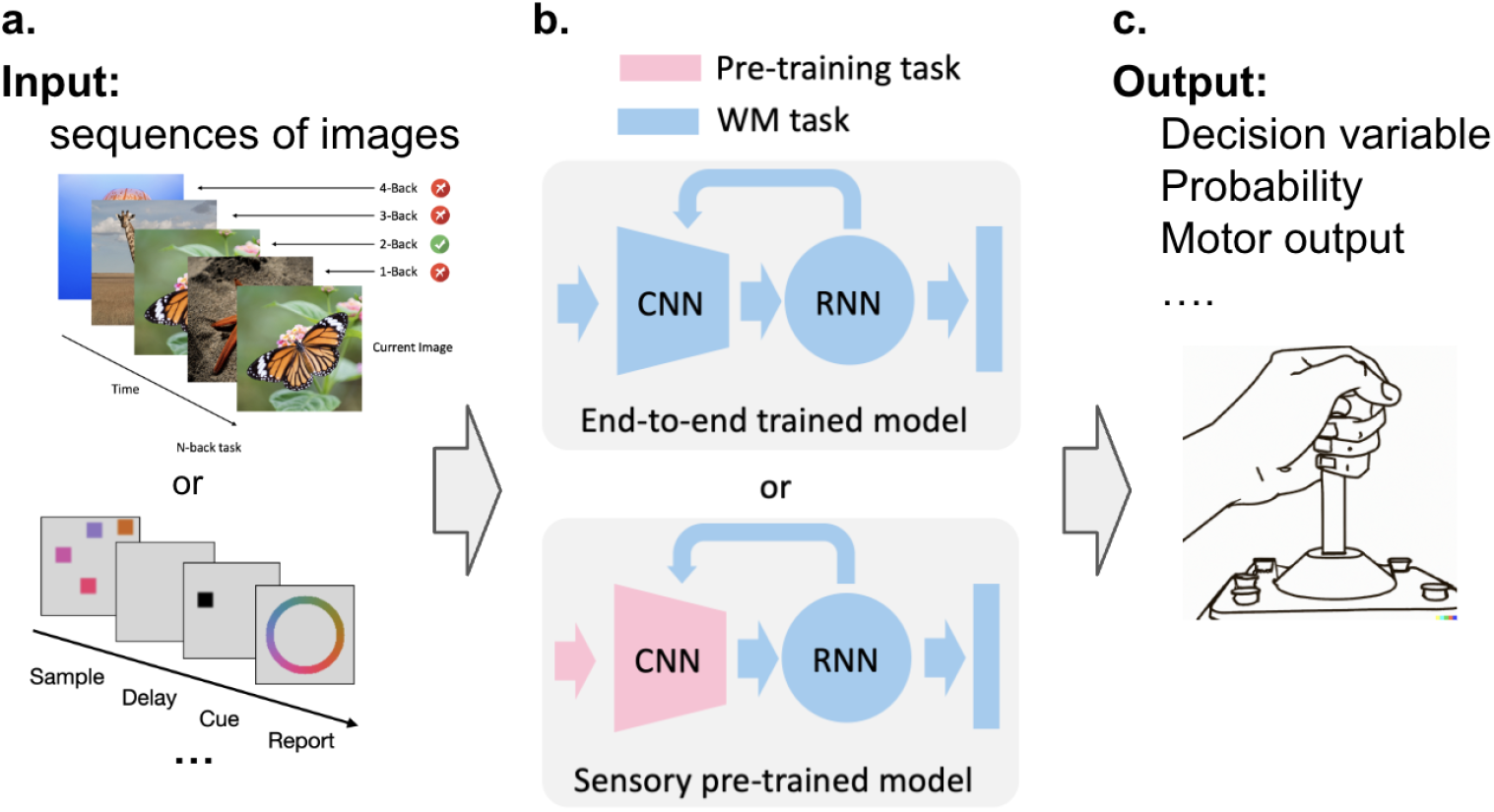
Image-computable sensory-cognitive models of visual working memory. (a) The inputs to our models are temporal sequences of images. Our image-computable model can be applied to arbitrary stimuli in raw pixel space, ranging from natural images (top) to abstract visual stimuli used in psychology or neuroscience experiments (bottom). (b) Schematics of the models we constructed. The models comprise two parts, the sensory system, and the cognitive/memory system. The sensory system is modeled by a CNN that receives an image as input and generates an output (a sensory embedding) at each time step. The cognitive/memory system is modeled by an RNN, which receives the sensory embedding and its own recurrent input at each time step to generate the behavioral output. The cognitive/memory system also generates top-down feature and spatial modulation to the sensory system [31]. The two-system model could be trained using different schemes. (Top) The entire model can be trained end-to-end on a working memory task. (Bottom) The sensory system can be pre-trained on other tasks and fixed before the cognitive/memory system is trained on a working memory task. The colors here indicate training objectives. (c) Our models generate a sequence of outputs, such as decision variables or motor outputs, depending on the tasks it is trained to perform.

### 2.2 Naturalistic pre-trained models exhibit human-like capacity limitation in change detection tasks

Seminal studies investigating the visual working memory capacity have employed the change detection paradigm [4, 13]. During the sample period of these tasks, human or animal subjects are presented with an array of stimuli, such as colored squares or oriented bars. Following a brief delay period, during which the stimuli disappear, the same array of stimuli is displayed again during the test period. However, unbeknownst to the subjects, some stimuli may change color or orientation. The subjects’ task is to report whether a change has occurred. Human performance in detecting changes rapidly declines as the number of stimuli to be remembered increases, a phenomenon known as the set size effect. This effect supports the notion of limited working memory capacity, as humans cannot hold an arbitrary number of stimuli in mind.

We trained our models to perform the change detection task using images with varying numbers of colored squares (Fig.2a). The models are trained through supervised learning to determine at the end of each trial whether a color change has occurred. When both the CNN and RNN are trained end-to-end on this task, our models exhibit super-human performance, achieving near-perfect accuracy regardless of the tested set size (Fig.2c). This finding suggests that the sensory-cognitive model architecture studied here possesses the capacity to perform the task almost flawlessly, surpassing human ability.

However, we observed strikingly different results when imposing naturalistic constraints on our models. Modern computational models of the ventral visual stream rely on convolutional neural networks (CNNs) trained on large-scale natural image tasks [27, 28]. After pre-training the sensory regions of our models with similar procedures and freezing the sensory region during change detection task training, our models’ performance closely aligns with that of humans [4]. Detection accuracy rapidly declines as set size increases, indicating a limited capacity (Fig.2c,d). We will further discuss the differences between end-to-end trained and sensory pre-trained models in subsequent sections.

In addition to the set size effect, our sensory pre-trained models match a wide range of experimental findings in change detection tasks. We analyzed the hit rate and false alarm rate of our model decisions, and found that as the set size increases the hit rate decreases and false alarm rate increases (Fig.2e), consistent with human data [37] (Fig.2f). We recorded our model’s decision confidence and performed Receiver Operating Characteristic (ROC) analysis, revealing smoothly increasing ROC curves within each set size (Fig.2g). This result is similar to human data [13], and is seen as a key support for the resource models (Fig.2h). In another experiment, we varied the color change magnitude in trials with a change, finding that the proportion of reporting a change increases with the change magnitude and the slope of the increase declines with the set size (Fig.2i), again mirroring human behavior [37] (Fig.2j).

These results indicate that naturalistic sensory pre-training imposes a strong and robust constraint on our models, leading them to exhibit human-like capacity limitations, aligning with a wide range of experimental findings.

**Fig. 2.**
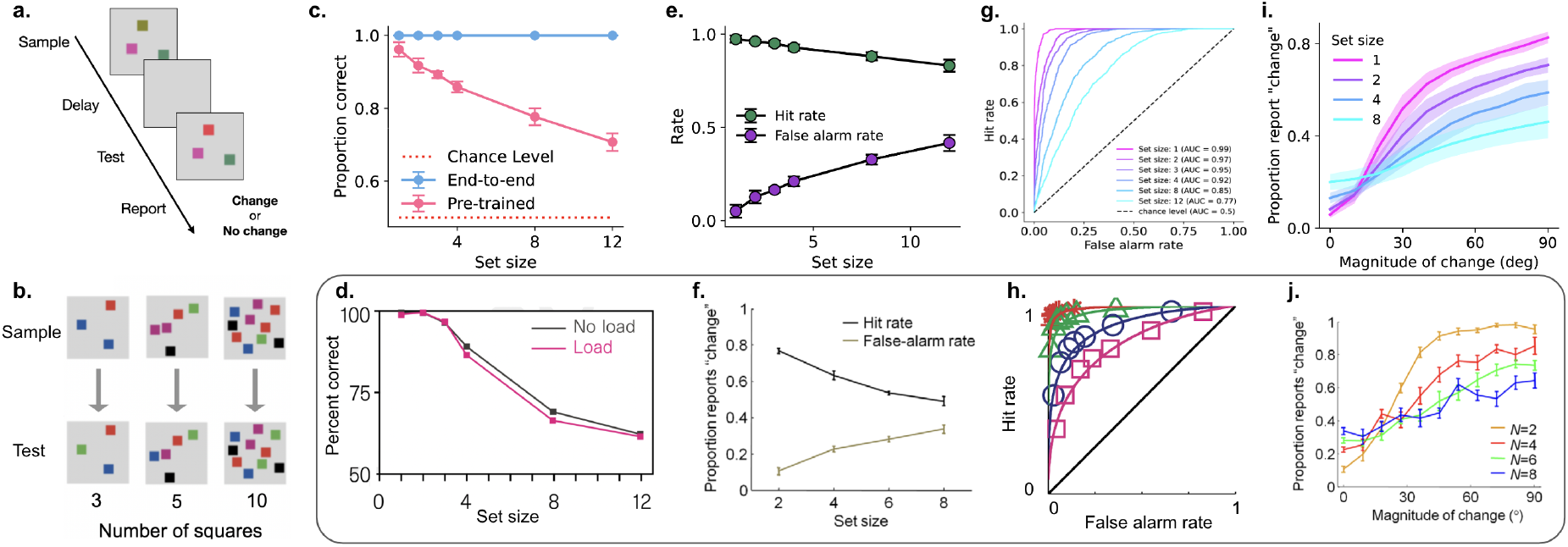
Naturalistic pre-trained models exhibit human-like capacity limitation in the change detection task. (a) A schematic of the change detection task. (b) Three example trials with different numbers of stimuli (set size 3, 5, 10). (c) The performance of end-to-end trained and sensory pre-trained models on change detection task with different set sizes. (d) Human performance on change detection task, from ref. [4]. (e) The hit rate and false alarm rate of model decisions in the change detection task at different set sizes. (f) the same analysis but for behavioral data, from ref. [37]. (g) ROC analysis of model decisions in change detection at different set sizes. (h) The same analysis, but for behavioral data, from ref. [13]. (i) The proportion of reporting a change in our model varies with the change magnitude at different set sizes. (j) The same analysis, but for behavioral data, from ref. [37].

### 2.3 Capacity limitation in delayed estimation tasks

In this section, we demonstrate that naturalistic pre-trained models exhibit human-like capacity limitations in another widely-studied task paradigm: delayed estimation tasks [5, 13]. These tasks provide a more detailed assessment of working memory content compared to the change detection task, which only requires binary decisions from subjects. In one example of delayed estimation tasks (Fig.3a), subjects observe an array of stimuli, such as colored squares, during the sample period. Following a brief delay, one stimulus location is cued, and subjects must report the color of the cued stimulus in a continuous space based on their retained memory. Experimental findings reveal that humans’ report errors increase rapidly as the number of stimuli (set size) increases [8], indicating a capacity limitation in working memory.

Consistent with our findings in change detection tasks, end-to-end trained models exhibit near-perfect performance with very low error, even when the set size is large (blue curve, Fig.3d). This again indicates that our model architecture possesses the capacity to perform the task far beyond human capabilities. In contrast, sensory pretrained models display rapidly increasing report error with set size, similar to human behavioral data [8] (Fig.3b-e). Although report precision decreases with set size in both end-to-end trained and sensory pre-trained models, only that of the sensory pretrained model can be accurately fit by a power-law function (Fig.3f), as seen in human data [6] (Fig.3g).

In addition to the set size effect, we found our sensory pre-trained model accounts for a wide range of other experimental findings in delayed estimation tasks, many of which have been essential to the debate between slot and resource models. The human report error distribution in delayed estimation tasks is characterized by a unique shape, distinct from the von Mises distribution, featuring a sharper peak and flatter tails [38]. Our model can also explain this unique shape (Fig.3b). When the error distribution of our models is fitted with a mixture of von Mises and uniform distributions, the residual between the distribution and the fit resembles human data [38] (Fig.3h,i).

In some versions of the delayed estimation task, a prioritizing cue appears before the sample period, indicating that one of the stimuli has a higher probability of being probed later. Humans exhibit enhanced memory precision for the cued item at the cost of precision for the uncued items [40], and this effect increases with cue validity. This is seen as support for a rational allocation of limited continuous resources for working memory. Our sensory pre-trained models demonstrate similar behavior [8] (Fig.3j,k). Our models also account for behavioral findings where subjects report their confidence during memory reports. In our sensory pre-trained model, when we require the model to report a confidence rating (see Methods), the error distribution widens when confidence is lower, even when the set size is fixed (Fig.3l), consistent with experimental findings [39] (Fig.3m).

Another well-known phenomenon in delayed estimation tasks is the feature binding error or swap error. Experimental subjects occasionally report the feature of a distractor stimulus instead of the cued stimulus [6]. The root of this swap error is not well understood and poses challenges for even the most recent visual working memory models, which often have overly simplified stimulus encoding assumptions. Our sensory pre-trained models exhibit swap errors (Fig.3n,o), opening the door for further investigation of the underlying mechanisms of this error.

**Fig. 3.**
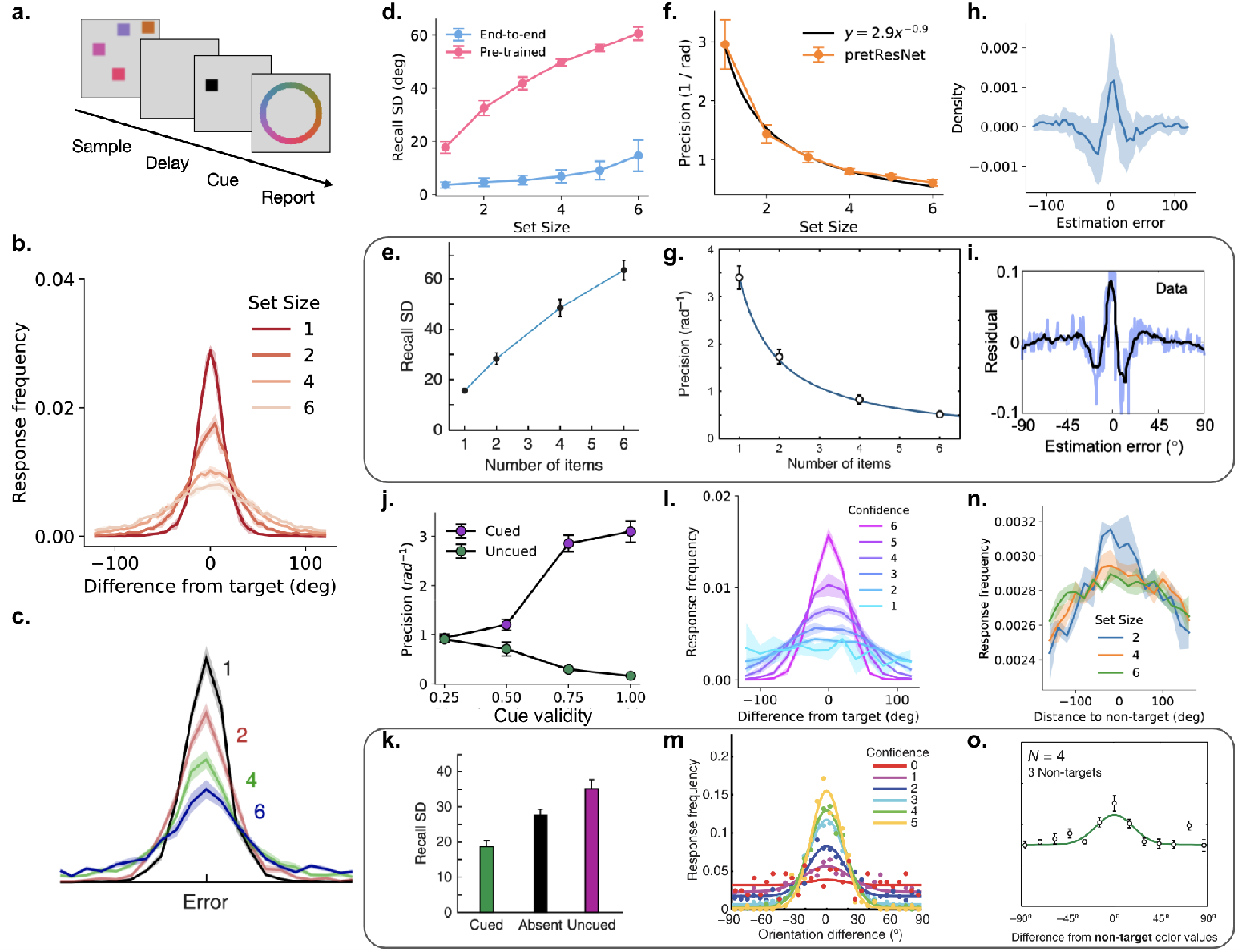
Naturalistic pre-trained models exhibit human-like capacity limitation in the delayed estimation task. (a) A schematic of the delayed estimation task. (b) The report error distribution with different set sizes from the the sensory pre-trained model. (c) The same analysis with behavioral data, from ref. [8]. (d) The standard deviation of the report error distribution (Recall SD) at different set sizes in sensory pre-trained and end-to-end trained models. The report error is measured as the angular difference between the reported color and ground-truth color in the color ring. (e) The same analysis with behavioral data, from ref. [8]. (f) The precision of report (reciprocal of the standard deviation of the report distribution) decreases with set size in the sensory pre-trained model. This trend can be fit by a power law function. (g) The same analysis with behavioral data, from ref. [6]. (h) The residual of report error distribution from the sensory pre-trained model. The residual is the report error distribution subtracted by a mixture of von Mises and uniform distribution fitted to the same distribution. (i) The same analysis with behavioral data, from ref. [38]. (j) The model report precision of an item cued or un-cued by a prospective prioritizing cue with different cue validity. (k) Analyses of behavioral data reveal that the cued stimulus have higher precision compared to uncued stimuli, from ref. [8]. (l) The report error distribution of the sensory pre-trained model grouped by different model confidence levels when the set size is fixed at 6. (m) The same analysis with behavioral data, from ref. [39]. (n) The report distribution of sensory pre-trained model centered around non-target distractors. (o) Non-target report distribution when the set size is 4 in behavioral data, from ref. [6].

In summary, our results demonstrate that naturalistic sensory pre-trained models account for a wide range of experimental findings in delayed estimation tasks.

### 2.4 Constraints other than naturalistic sensory pre-training do not lead to human-like capacity limitation

In sensory pre-trained models, the connection weights in the sensory neural network are established during pre-training and remain fixed when trained on working memory tasks. Sensory pre-training can be viewed as a form of constraint imposed on end-to-end trained models. It is possible that introducing other constraints to end-to-end trained models could also result in human-like capacity limitations. Therefore, we investigated the effect of other constraints, such as limiting the model size and adding sensory or memory noise.

In the change detection task, we reduced the number of channels in the CNN or the number of hidden units in the RNN of the end-to-end trained models. We found these size constraints have modest impact on the performance of the end-to-end trained models. End-to-end models still perform much better than humans and have no human-like set size effect even with a significantly smaller model size (Fig.4b). This result suggests that the model architecture we used has ample capacity to perform the task almost perfectly when trained end-to-end on the task.

Sensory or memory noise is an important factor that results in capacity limitations in many previous models of working memory [13, 14]. To investigate whether noise in our model can also lead to human-like capacity limitation, we added various levels of sensory or memory noise to end-to-end models during evaluation on the change detection task (Fig.4c,d) or the delayed estimation task (Fig.4f,g). The sensory noise is added to each input image, and the memory noise is added to the recurrent units of the RNN. When sensory noise is added to the model, performance on the change detection task decreases and the report error increases almost uniformly regardless of set size (Fig.4c,f). When memory noise is added, the performance on both tasks still show strong uniform decreases. Although we see signs of set size effect in these conditions (Fig.4d,g), the performance on these tasks does not resemble behavioral data seen in Fig.2 and Fig.3.

In summary, human-like capacity limitation cannot be easily obtained by adding many other constraints to our end-to-end trained models, such as limited model size or sensory and memory noise. Our results suggest that naturalistic pre-training may be an essential constraint for models to exhibit human-like capacity limitation.

**Fig. 4.**
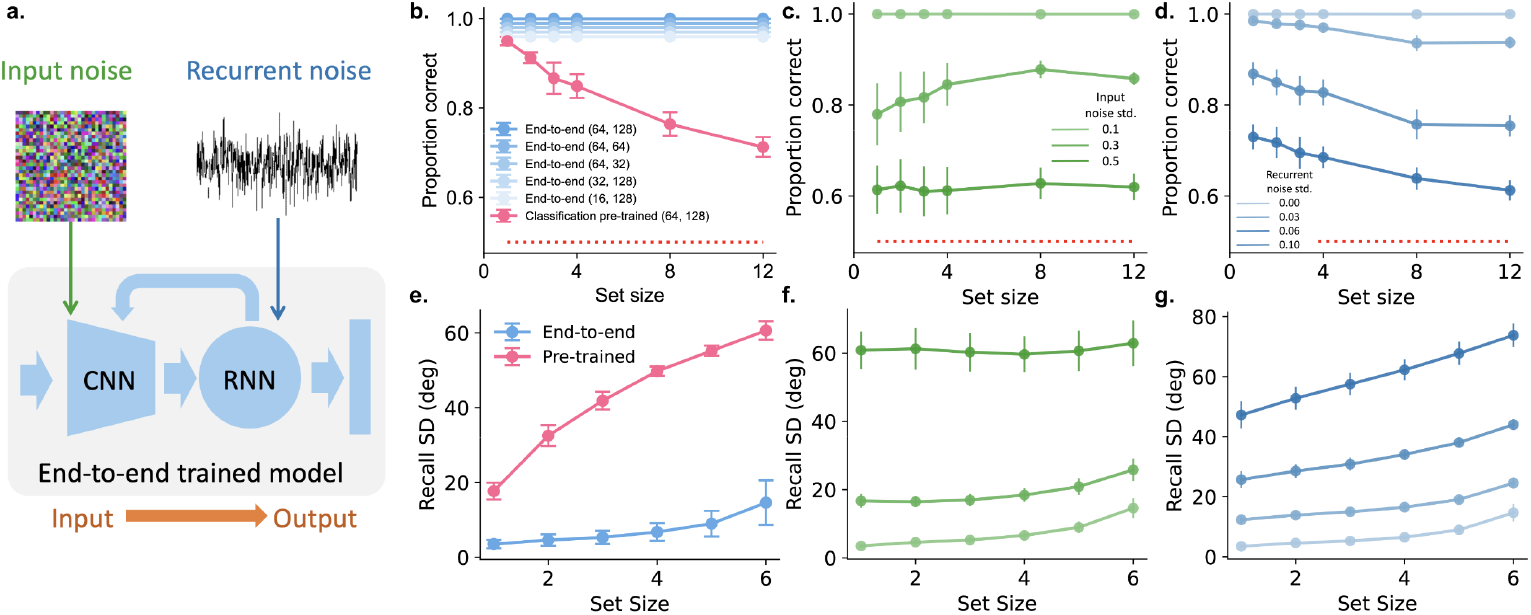
Limiting model size or adding noise to model do not lead to human-like capacity limitation in end-to-end trained models. (a) Two different ways to add noise to the model. Input noise is applied to the images as inputs to the CNN. Recurrent noise is applied to recurrent units of the RNN. (b) Change detection performance of end-to-end trained models with different CNN or RNN sizes. Number inside parentheses indicate sizes of the model: (number of CNN channels, number of RNN hidden units). End-to-end models are slightly offset for visualization purpose. (c) Change detection performance using end-to-end trained models with different levels of input (sensory) noise. Each curve indicates the performance at one noise level. Darker colors indicate higher noise level (from light to dark, standard deviation of the random normal noise: 0.1, 0.3, 0.5). Inputs noise are added to the input images. The open error bars here indicate standard errors, and closed error bars on other figures indicate standard deviation (SD). (d) The same as (c). but with different levels of recurrent (memory) noise added to the RNN. Darker colors indicate higher noise level (from light to dark, standard deviation of the random normal noise: 0, 0.03, 0.06, 0.1). (e-g) The same as in (b-d), but for the delayed estimation task. The curves here show the standard deviation (Recall SD) of the report error distribution at different set sizes.

### 2.5 Robustness to hyperparameters and model architectures

The size of neural network models is typically critical in determining their capacity. It is possible that the human-like set size effect in our sensory pre-trained models trivially results from specific model size hyperparameters. To investigate whether the observed set size effect in sensory pre-trained models is robust to hyperparameters, we varied the width of the CNNs or the number of units in the RNNs in these models. We discovered that the set size effect in naturalistic pre-trained models is robust to the model size once the model size exceeds a small threshold (Fig.5a). We further examined models with other CNN architectures pre-trained with images from ImageNet [34] with a larger input size (224×224 pixels). These models also demonstrate a decrease in performance when we increase the set size, although the magnitude of the decline is smaller (Fig.5b). The performance of models with randomly initialized versions of these CNNs does not show an apparent set size effect (Fig.5c).

We further investigated other pre-training paradigms for the sensory regions. Contrastive self-supervised learning can be used to train CNNs using unlabeled images, and has been shown to result in CNNs that capture visual responses of monkeys [27] and mice [41]. Our models also exhibit the set size effect when the CNNs are pre-trained on natural images with contrastive learning (Fig.5d). However, when the sensory region is pre-trained to recognize handwritten digits or merely randomly initialized, the models’ performance is much closer to chance performance, and the set size effect is weaker (Fig.5d).

**Fig. 5.**
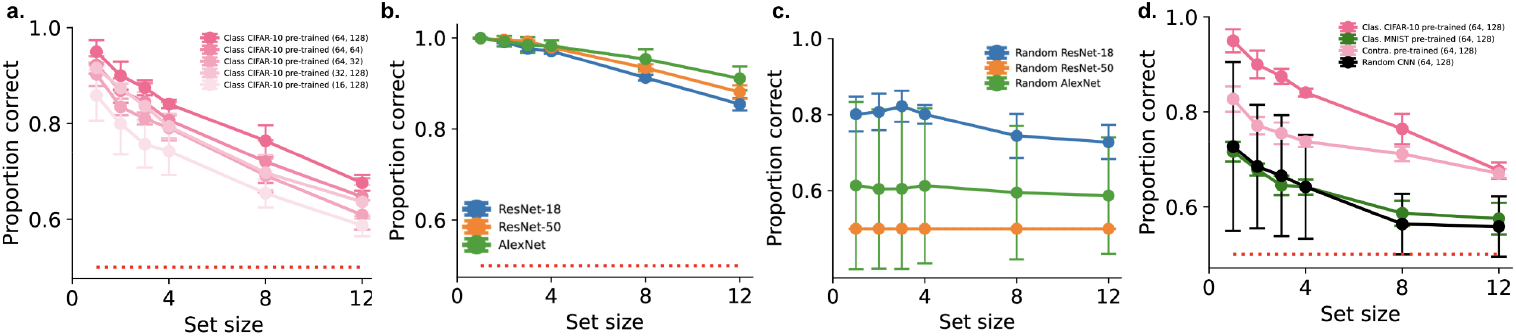
Robustness of human-like capacity limitation to hyperparameters and different model architectures. (a) Change detection performance of sensory pre-trained models with different CNN width and RNN size. Number inside parentheses indicate sizes of the model: (number of CNN channels, number of RNN hidden units). (b) Change detection performance of models with larger CNNs that receives 224 by 224 sized images and pre-trained on ImageNet. (c) The same as (b), but for models with randomly initialized CNN weights. (d) Change detection performance of models with different sensory pre-training paradigms.

These results indicate that each of the separate factors—model architecture, training objective, and training data—is in itself insufficient to explain the human-like set size effect. The human-like set size effect only occurs when we combine pre-training the convolutional network on naturalistic images with the appropriate objective.

### 2.6 Bottom-up capacity limitation from sensory constraints

Behavioral models of visual working memory capacity frequently depend on abstract constructs such as slots, resources, or noise. However, the neural substrates corresponding to these constructs remain largely unidentified. In this paper, we offer a preliminary analysis of neural activity in our models, aiming to potentially illuminate the neural mechanisms underlying capacity limitations.

A commonly used marker for working memory-related activity in humans is contralateral-delay activity. Studies suggest that the amplitude of this delay activity correlates with the number of items remembered, reaching a plateau when more than three or four stimuli are presented in a single trial [42]. Experiments analyzing fMRI data have demonstrated that BOLD signals initially increase and then plateau as more items are presented [43]. We recorded average neural activation in the CNNs of our sensory pre-trained models and found that neural activation in the middle layer of the sensory CNN exhibits similar trends to those observed in neural data (Fig.6a,b).

Neurophysiological data obtained from monkeys performing working memory tasks imply that interference between representations of multiple items, occurring in a bottom-up manner, may serve as one potential mechanism underlying capacity limitation [44]. To further explore the cause of capacity limitation in our sensory pre-trained models, we examined the detectability of changes at various layers within our models. Specifically, we computed the neural activity of our model in response to both the sample stimulus (an array of colored squares) and the test stimulus. We then trained linear decoders to determine whether a change exists between the sample and test stimuli, using neural activities from different model layers. Our findings reveal that in the early sensory layers, neural activity retains sufficient information for the linear decoder to nearly perfectly detect potential changes, irrespective of set size. As we progress through the sensory hierarchy towards the cognitive region, information about change or no-change gradually diminishes (Fig.6c). These results suggest that capacity limitation in our model arises in a bottom-up manner, consistent with electrophysiological observations.

To gain a deeper understanding of the nature of the interference, we conducted a state space analysis on the neural activities. Our analysis revealed that as more distractors are added, the neural manifold encoding a single color patch shrinks (Fig.6d-g). This implies that interference between multiple items results in the shrinkage of representation for each item, potentially underpinning the increased error observed when multiple items are retained in memory.

**Fig. 6.**
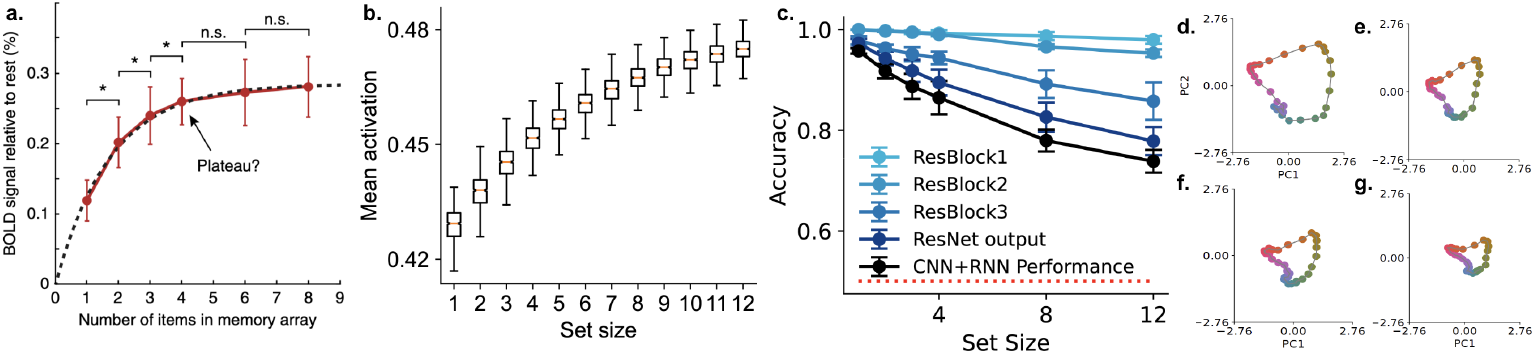
Neural mechanisms of capacity limitations in our models. (a) fMRI BOLD signal amplitude in parietal regions when different number of stimuli are presented, from ref. [8]. (b) Overall activation in the second convolutional layers of the CNN in our model when presented with different number of stimuli. (c) Accuracy of decoding a change or no-change in the change detection task using neural activities at different layers in our model. ResBlock 1,2,3 mean the first, second, and third convolutional blocks in the CNN, ResNet means the CNN output to the RNN. (d-g) The neural state space encoding for one color patch at the presence of 0, 1, 2, and 3 distractors.

### 2.7 Capacity limitation with sequentially presented stimuli

In most working memory tasks involving multiple items, the items can be presented either simultaneously or sequentially. Our previous results were obtained when multiple items (colored squares) were presented simultaneously. We found that capacity limitation arises in a bottom-up fashion through interference building up already in the sensory layers. However, when items are presented sequentially, no sensory interference exists in the feedforward CNN we studied. It remains unclear whether capacity limitation may still be observed in our models. Consequently, we explored a version of the delayed estimation task in which stimuli were presented sequentially (Fig.7a).

We found that the report error of the sensory pre-trained models increases rapidly with set size (Fig.7b), demonstrating a form of set-size effect and, therefore, capacity limitation. However, unlike the case of simultaneous presentation, the capacity limitation is only observed when the number of units in the RNN is relatively low. As more units are added to the RNN, the working memory capacity of the model easily surpasses human capacity and appears to increase without bound (Fig.7c). Another prominent effect in sequentially presented delayed estimation tasks is the recency effect, in which the last stimulus in a sequence is remembered with higher accuracy than other stimuli when probed. However, an analysis of our model’s report error distribution does not reveal such a recency effect (Fig.7d).

**Fig. 7.**
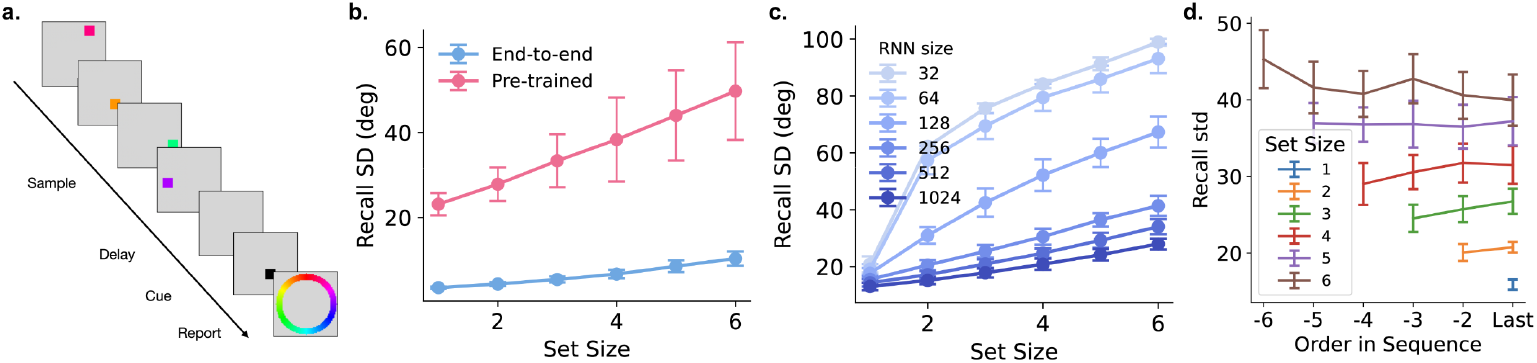
Capacity limitation in working memory tasks with sequentially presented stimuli. (a) A schematic of the delayed estimation task where stimuli are presented sequentially. (b) The standard deviation of the report error distribution (recall SD) of a sensory pre-trained model and an end-to-end trained model in this task. (c) The performance of sensory pre-trained models with varying numbers of hidden units in the RNN. (d) The recall SD of stimuli at different rank orders in the stimulus sequence, across various set sizes.

These results suggest that, in tasks with sequentially presented stimuli, constraining the RNN may be necessary to explain the observed capacity limitation in behavioral data. We can select a particular RNN size (*∼* 100) that results in the set size effect seen in behavioral data; however, this is not a realistic constraint, as millions of parietal and prefrontal neurons in mammalian brains may be involved in working memory tasks. Given the results in sequentially presented tasks, further investigation into specific constraints on model architecture or training objectives, particularly focusing on the cognitive/memory region, is needed to fully explain the nature of capacity limitations.

## 3 Discussion

In this study, we developed a new kind of computational model for visual working memory by combining CNNs and RNNs. Our models consider more realistic sensory encoding and can process raw sensory stimuli in pixel space. Without explicit assumptions of slots or resources, we discovered capacity limitations resulting from pre-training the sensory region of our model on naturalistic tasks using real-world images. These sensory pre-trained models account for a wide range of experimental findings in change detection and delayed estimation tasks. Additionally, our neural network models enable us to analyze neural activities to elucidate the neural mechanisms underlying capacity limitations. We found that capacity limitations in our model occur in a bottom-up fashion and are likely due to interference between multiple items in the sensory system. Our modeling results demonstrate that sensory pre-training imposes sufficient constraints on the model to account for a broad range of experimental findings.

Our results suggest two broader implications. First, even when studying memory or other high-level cognitive tasks, it may be crucial to construct models that account for realistic sensory processing rather than employing oversimplified stimulus representations, as many memory phenomena are likely tightly linked to the specific sensory domain. Second, when building neural network models of the brain, it is essential to consider different neural systems with potentially distinct objectives and architectures instead of a monolithic model trained end-to-end on a single objective.

The importance of realistic sensory processing in working memory models has been explored in a limited fashion recently. Schurgin and colleagues have pointed out the importance of considering the perceptual similarity in the space of colors or faces used for working memory task [9]. Hedayati and colleagues proposed a variational autoencoder model formed by CNNs and trained on images as a computational model for working memory [45]. Using CNNs allow the model to remember familiar items using neural representation from higher layers of the visual hierarchy while remembering novel items using lower-level representation. However, the rich and detailed impact of sensory processing on visual working memory capacity limitation has not been characterized before.

Phenomenological models, such as the slot-based [7], resource-based [8], and signal detection models [9, 13], have demonstrated superior quantitative fits for human behavior in working memory experiments. In contrast, our model only provides a qualitative match to human behavior. It remains to be seen whether the network can yield a quantitatively precise match by adjusting selected parameters, such as noise level or model size.

Despite this limitation, our model presents several fundamental advantages. In addition to offering potential neural mechanistic accounts, as expected of a neural network model, our model gives a principled and functionally grounded account for the working memory capacity limitation. Consider the recent model based on signal detection theory [9], which provides a superior and parsimonious fit to human error patterns in the delayed estimation task. This theory explains the set size effect by fitting different memory strength parameters to various set sizes. However, it needs to explain why memory strength should decay rapidly, rather than more gradually, with increasing set size. Our model proposes that the strong working memory limitation arises from a fundamental “mismatch” between the objectives of the working memory and visual system and the tasks employed to assess working memory.

From an evolutionary perspective, the working memory system’s purpose is presumably to temporarily store and manipulate critical information that informs future actions, both external (movement) and internal (attention). The essential information to be stored and manipulated is likely in the form of complex objects or concepts rather than the colors of arbitrary patches. Consistent with this hypothesis, colors are better remembered when they are part of meaningful objects [46], and human’s working memory performance is higher for real-world objects [25].

Although our modeling results align qualitatively with a wide range of experimental findings, some of our results do not provide a detailed match to behavioral observations. For instance, in the original change detection study, human performance remains high until it suddenly drops at 3-4 items [4]. Our modeling results do not reproduce this sudden drop at a specific set size. The detailed performance profile might depend on the particular experimental conditions of the original study, such as the colors used, the size of the color patches, or the extent of the color change. Our reproduction of the task aims to capture the essential features of those tasks, but the experimental conditions are not identical. Instead of seeking precise quantitative matches to experimental data, our objective here is to identify parsimonious design principles that lead to a wide range of qualitative matches to data.

Contrasting our simultaneous and sequential presentation results reveals that the root of capacity limitation may be multi-faceted. In a simultaneous presentation task, the capacity limitation likely arises from the interference of multiple stimuli in the sensory system. However, in sequential presentation experiments, humans or animals can encode each item without distractors but still exhibit limited working memory capacity. In other experiments, even when the stimuli are presented simultaneously, subjects have ample time to focus on each item individually but still exhibit capacity limitations. Although we observed set size in sequential experiments by constraining RNN size to a few dozens of neurons, it is still puzzling why humans or other animals have a small capacity while having billions of neurons. Future works need to be done to gain a more principled understanding of capacity limitation in sequential presentation. As for what causes capacity limitation in sequential tasks, we have two speculations. First, more constraints on the RNN memory model may be required. For example, the RNN may need to be pre-trained on ethologically motivated tasks as well [47], such as video predictions. Or some other regularization on the activation or connection weights of the RNN may be needed [32]. Second, we may consider alternative model architectures. Experimental evidence from human neuroimaging and non-human physiological studies suggests that working memory is stored in a distributed fashion across cognitive regions and the sensory hierarchy used to process ongoing sensory information [48, 49]. Consequently, interferences may still exist within the sensory hierarchy even when stimuli are encoded sequentially. Sensory-cognitive networks that support working memory with a feedback loop between the visual and cognitive systems may be essential to explore these questions. Besides long-range feed-back loops, the visual system may need local recurrent processing, modeled through convolutional recurrent neural networks [50].

## Methods

### Code sharing

The code for model, task, and data analysis will be made available at: https://github.com/metaconsciousgroup/multisyswm.

### Neural network models

Our models primarily consist of a Convolutional Neural Network (CNN) and a Recurrent Neural Network (RNN). The CNN processes raw images as input, and the penultimate layer serves as input to the RNN. The behavioral output is readout from the hidden units of the RNN. In some models, the RNN provides top-down modulation to the CNN. The entire model receives a sequence of images as input and generate a corresponding temporally dependent sequence of output simultaneously. In the change detection task, the output is a score variable indicating a binary decision of change or no-change. In the delayed estimation task, the output are two numbers indicating the horizontal and vertical coordinate of the reported location. When the confidence is needed to be reported, models with an additional output indicating the standard deviation of the reported position is used. For simplicity, we assume the horizontal and vertical coordinate have the same standard deviation. All models are implemented using the PyTorch library.

In most of our experiments, we employed a 20-layer ResNet convolutional neural network, following the design in [51]. This network architecture was designed to process 32 *×*32 sized images, such as those from the CIFAR10 dataset. Our specific implementation is adapted from the PyTorch library implementation found at: https://github.com/pytorch/vision/blob/main/torchvision/models/resnet.py. For Fig. 5c, we utilized pre-trained ResNet-18 or ResNet-50 to process larger images with dimensions of 224 *×*224. These models are trained on the ImageNet dataset. The weights of these models are downloaded from the PyTorch library.

The RNN component of our model is based on the Continuous Time RNN (CT-RNN) architecture. This kind of recurrent neural network is based on the differential equations used to model spike rate in biological neurons and are commonly used in neuroscience to better capture realistic neural dynamics. The update rule at each time step is:

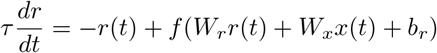

In the discrete form, it is:

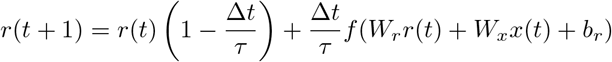

Here *r*(*t*) is a vector of the firing rate of the hidden units at time step *t, W*_*r*_ is the hidden weight matrix, *W*_*x*_ is the input weight matrix, *b*_*r*_ is the bias. ∆*t* and *τ* are time constants. Here we used 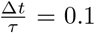 is the non-linear activation function. Here we used tanh nonlinearity.

Top-down modulation is incorporated into the model using the Convolutional Block Attention Module (CBAM) [31] approach. At each time step, the RNN generate a feature modulation vector and a spatial modulation vector that are then applied to the CNN activation in the next time step. The modulation is applied to the second convolutional layer in the last ResNet block of each of the 3 layer. The CNN activation is multiplied by the modulation vector and then passed to the next layer.

### Tasks

All of our tasks uses a sequences of images as input, intended to mimic the tasks used in human or animal experiments.

Change detection task. A single trial of the change detection task has three periods, sample, delay, and test. In the sample period, the stimulus is an image with several colored patches on a gray background. The image has the dimension of 32 *×*32 and each color patch has the dimension of 5*×* 5. The positions of the colored patched are sampled randomly while avoiding overlap between different patches. The colors of these patches are sampled randomly from a set of 6 distinct colors, or sampled from a continuous color ring when we need to vary the magnitude of the color change. The color ring is in the CIE L*a*b color space, centered at L = 54, a = 21.5, and b = 11.5, with a radius of 49. In the delayed period, the color patches disappear and a blank gray image is used as the input. In the test period, the same image in the sample period is used as input, but 50% of the time, the color of one of the patch could be changed to another randomly sampled color. The model need to remember the sample stimulus and use that information to make a decision of on whether there is a change of color or not.

Delayed estimation task. A single trial of the delayed estimation has three periods, sample, delay, cue/report. In the sample period, similar to the change detection task, the stimulus is an image with several randomly sampled colored patches on a gray background. The colors are sampled from the color ring mentioned before. The patches disappear in the delay period. In the cue/report period, a black patch appears in the same location as one of the colored patches shown during the sample period, indicating the model need to report the color of that cued patch. The model generate a response to report the color of the cued patch on a continuous color ring. The report output is the horizontal and vertical coordinate of the color on the color ring. The ground-truth color coordinate in the report ring is the same as the ring used to sample the colors. Delayed estimation task with prioritizing cues. This task is similar to the above-mentioned delayed estimation task, except that an additional cue appears before the sample period, indicating that one of the squares is more likely to be probed later.

Delayed estimation task with sequential presentation. This task is similar to the above-mentioned delayed estimation task, except that there is a longer sample period. Each colored square is presented one-by-one to the model instead of being presented simultaneous.

### Model Training

Models are trained with gradient decent with Adam optimizer implemented by the PyTorch library.

Sensory Pre-training. For models that involves sensory pre-training, the CNN part of the model is first trained on a sensory task. In classification pre-training, the 20 layer ResNet is trained on image classification on the CIFAR10 dataset, which contains 10 categories of natural images. For Fig 5c, we used ImageNet pre-trained ResNet-18 and ResNet-50 downloaded from the Torchvision library. In contrastive pre-training, we follow the contrastive pre-training pipeline in [36] and trained the CNN using images from the CIFAR10 dataset. The code for contrastive pre-training is adapted from https://github.com/sthalles/SimCLR.

Supervised task training. After sensory pre-training, we drop the last layer of the CNN and connect the penultimate layer to the RNN. Then the entire model, including parameters used to generate attention modulation, is trained on the working memory task using supervised learning to minimize the error between output of the model and ground truth. Separate models are trained for the change detection task and the delayed estimation task. In the change detection task, the loss is the Binary Cross Entropy between the target and the input probabilities. In the delayed estimation task, the loss is the mean squared error between the model output and the ground truth.

In sensory pre-trained models, the weights of CNN is fixed during supervised task training. In end-to-end trained models, the weights of CNN are randomly initialized and are trained with the RNN together on the task.

### Analysis of Behavioral and Neural Data

Change detection task. For ROC analysis in the change detection task, the model output score of the binary decision is recorded and used to perform the analysis. In the delayed estimation task, we recorded the model output error and fitted a mixture of von Mises and uniform distribution to the error distribution, as in [38]. The residual is calculated in the following ways. First, we pool all the error at different size to generate an overall report error distribution. Then, we subtract the best fitting mixture from the report error distribution.

## Acknowledgement

We thank Timothy Buschman, Nicholas Watters, Wei ji Ma, Samuel Gershman, Christopher Bates, Aran Nayebi, Albert Compte, and Ali Hummos for many helpful discussions and comments.

## Notes

### Competing Interest Statement

The authors have declared no competing interest.

